# The effect of the thin body ideal in a media-naïve population

**DOI:** 10.1101/176107

**Authors:** Jean-Luc Jucker, Tracey Thornborrow, Martin J. Tovee, Lynda G. Boothroyd

## Abstract

The thin ideal is the western concept of an ideally slim or underweight female body^1^, and its omnipresence in the mass media has a negative impact on women’s health^2-5^. Media consumption is associated with a drive for thinness, body dissatisfaction, low self-esteem, and disordered eating in women of western and/or industrialised societies^4^. Furthermore, cross-cultural research suggests that the media have similar effects when they are introduced into non-western or non-industrialised societies^2,6,7^. No study, however, has attempted to induce a change in female body size ideals in a population that is not exposed to the thin ideal and that has currently no access to the media. Here we show experimentally that a short exposure to the thin ideal can change body size ideals in a media-naïve population. 80 rural Nicaraguan men and women with very low to non-existent media access created their ideal female body before and after seeing photographs of either thin or plus size fashion models. Analyses revealed a significant interaction between time and group, meaning that exposure to media images shifted the subjects’ ideal female body size. We discuss problems posed by the pervasiveness of the thin body ideal in the context of the global obesity pandemic.

---

Up to 30-50% of girls and women from western or industrialised countries have negative body image^8,9^, which contributes to widespread psychopathologies such as low self-esteem, depression, and disordered eating^10,11^. One key sociocultural contributor to body dissatisfaction is the thin body ideal and its omnipresence in the mass media^1,3,5,12-14^. For example, across 77 correlational and experimental studies, media exposure was associated with a stronger internalisation of the thin body ideal, higher body dissatisfaction, and higher eating disorder symptomatology^4^. Furthermore, cross-cultural research has shown that populations with limited access to the media have a higher female body size ideal^7,15-17^, and that the introduction of television in previously media-naïve populations is accompanied by an increase in body dissatisfaction and pathological eating attitudes^2,6,7,14^.

Although past research suggests that the media impact female body size ideals and so have a negative effect on women’s health, it suffers from two important limitations. First, experimental research has never shown that the thin ideal and the media can induce a change in body size ideals in a non-WEIRD (western, educated, industrialised, rich, and democratic)^18^, media-naïve population. Instead researchers have used western, industrialised subjects in countries where the thin body ideal is already omnipresent in the media and where poor body image is already widespread^19-21^. Second, all cross-cultural research using non-western or non-industrialised subjects has been correlational, cross-sectional, or pseudo-longitudinal, and has therefore failed to establish causation between exposure to the thin ideal and a decrease in female body size ideals.

Here we describe the first study to experimentally induce a shift in female body size preferences using thin ideal stimuli in a non-western media-naïve population. To do so, 80 male and female subjects were drawn from two Nicaraguan villages with no access to grid electricity and thus very low to non-existent access to visual media (see Table 1 for sample characteristics). They were tested on their ideal female body size (hereafter ‘ideal body task’), before and after seeing photographs of either thin or plus size fashion models (hereafter ‘manipulation’). In the ideal body task, subjects used computer-generated female bodies to create their ideal female body based on multiple different exemplars or ‘starter bodies’ (Figure 1). They could do so by increasing or decreasing the size of these starter bodies within a range equivalent to BMI 15-40. The subjects created 4 bodies at pre-test, and 4 bodies (using different starter bodies) at post-test, with the order of starter bodies (pre vs post-test) counterbalanced across subjects.

**Table 1:** Sample Characteristics.

|  |  |
| --- | --- |
| Valid N | 71 |
| % Female | 49.29 |
| Age (years) | 30.42 (12.95) |
| Body mass index | 24.63 (4.98) |
| **Dieting** |  |
| % Not dieting | 69.01 |
| % Decrease weight | 18.30 |
| % Increase weight | 12.67 |
| Earnings (\$/year) | 1,230.06 (2,356.20) |
| Education (years) | 5.35 (4.80) |
| **Ethnicity** |  |
| % Garifuna | 29.57 |
| % Mestizo | 56.33 |
| % Mixed or other | 14.08 |
| **TV access** |  |
| % In own house | 8.45 |
| % In neighbour's house | 16.90 |
| % No access | 74.64 |
| **TV consumption** |  |
| TV consumption in last 7 days (hours) | 1.55 (3.14) |
| **TV type** |  |
| % TV set + DVD player only | 15.49 |
| % Satellite TV | 12.67 |
| % No TV | 71.83 |
| Time since last meal (hours) | 3.57 (1.84) |
| **Travel** |  |
| Last trip to other village (days) | 234.94 (859.03) |
| Duration of stay (days) | 4.25 (12.01) |
| Daily TV consumption (hours) | 0.77 (1.25) |

*Note.* 'TV access' refers to access to any type of television, not necessarily satellite television; 'TV type' specifies which type of television the subjects had access to. Further, 'TV consumption in last 7 days (hours)' includes any TV watched (and any type of TV watched) while travelling in the last 7 days.

**Figure 1.**
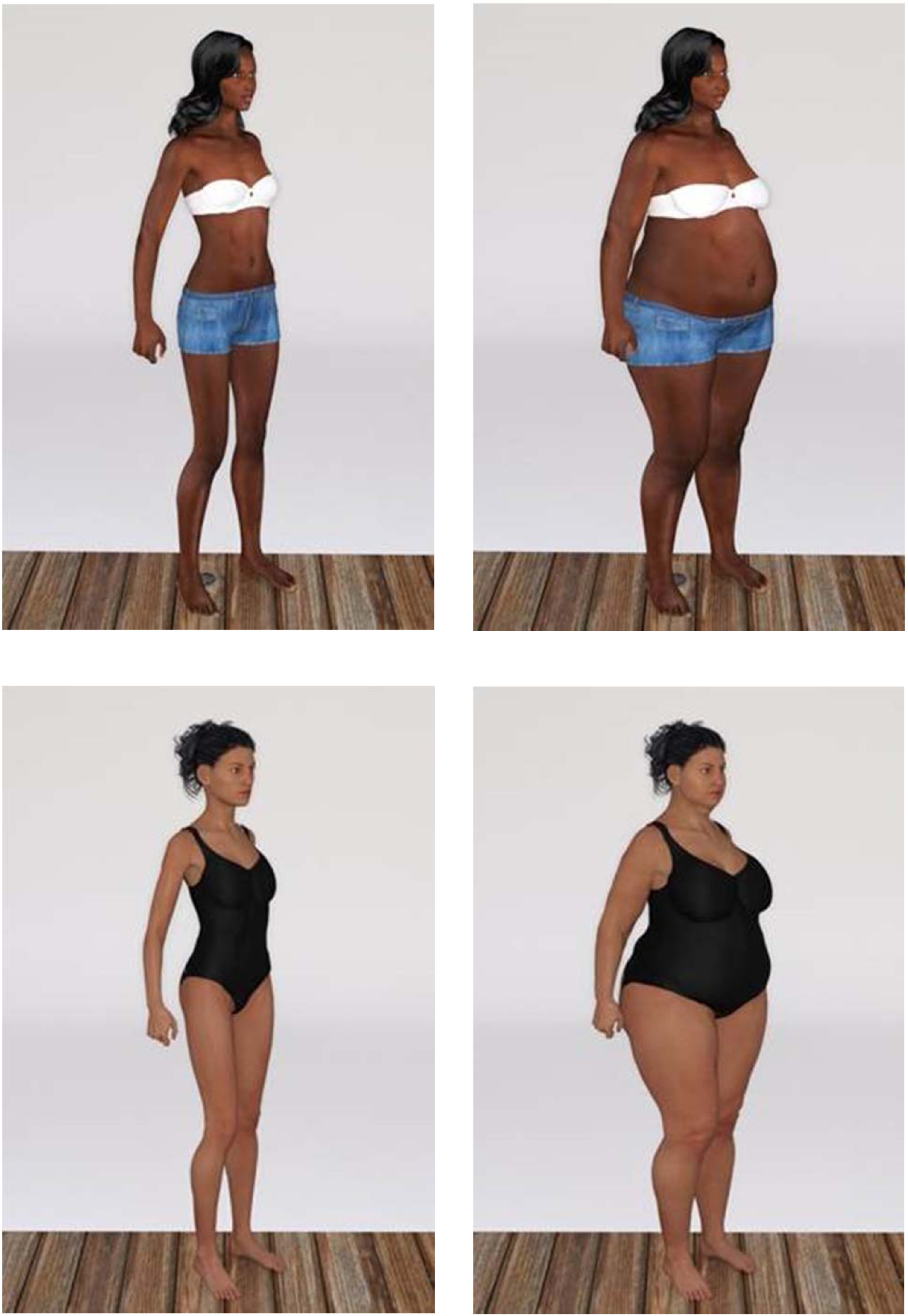
Sample starter bodies (minimum and maximum size)

In the manipulation phase, subjects were exposed to a series of photographs of either thin or plus size fashion models (Figure 2). The photographs were taken from western retailer’s catalogues or popular women’s magazine, and depicted models wearing clothes ranging from UK size 4-6 (thin models) to 16-28 (plus size models). The models varied in age, ethnic background, and clothing, with these characteristics being represented equally in both experimental groups. To ensure visual engagement with manipulation stimuli, the photographs were presented in pairs and the subjects were instructed to choose their favourite model out of each pair.

**Figure 2.**
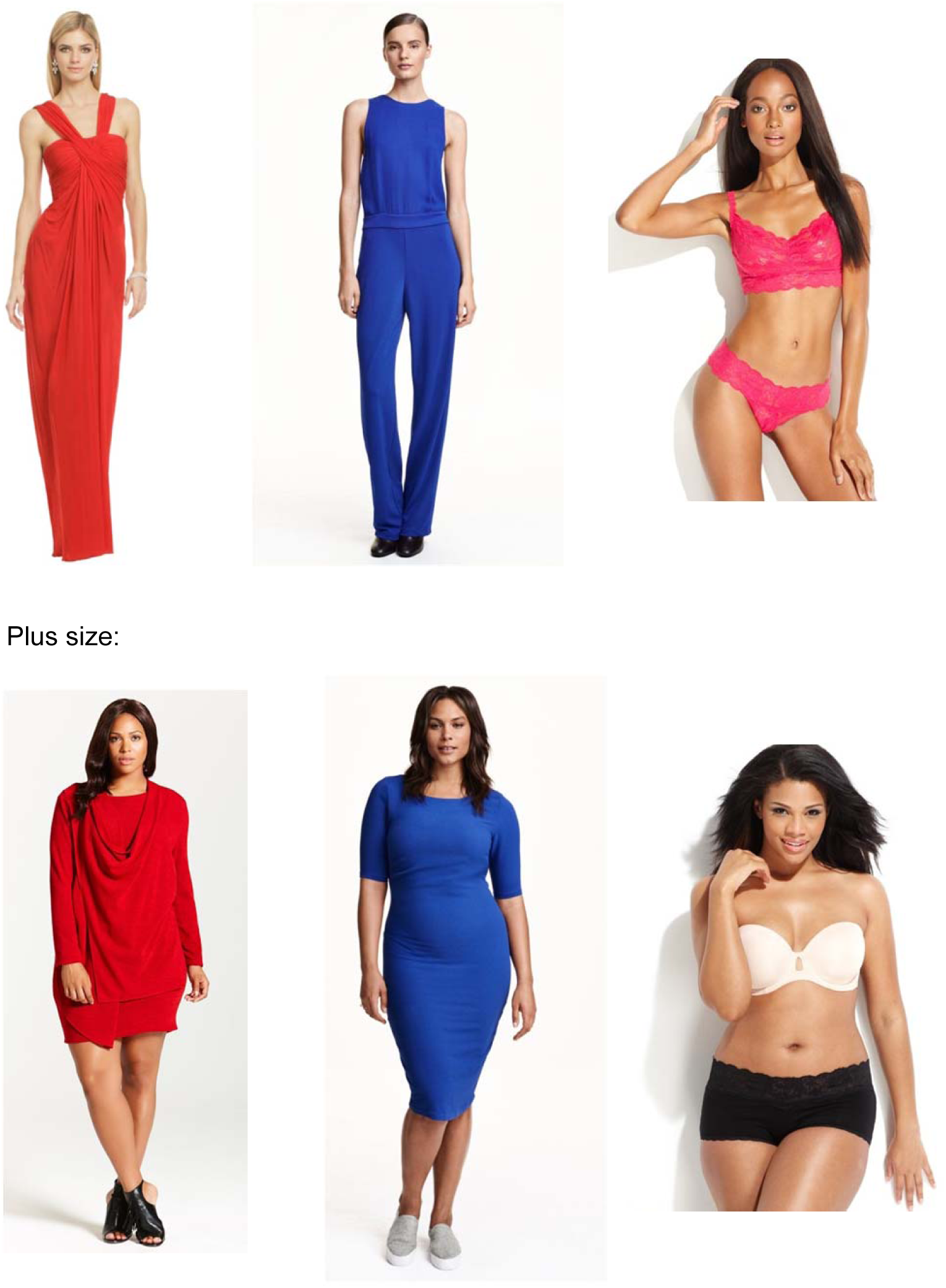
Sample thin size and plus size models.

Preliminary analyses showed that the two experimental groups were matched on all variables measured in this study, including age, BMI, dieting, earnings, education, ethnicity, television access and television consumption, time since last meal, sex, and travel (*F*s < 1.955, *p*s > .167). To determine whether time and group interacted, a random intercept model was constructed with subject as the higher level and body/trial as the lower level unit. Initial variance partitioning (Model I; Table 2) indicated that 52.8% of variance operated at the level of the body identity/trial, while the remainder operated at the level of the subject. Time (pre-test vs. post-test) was entered as a body-level covariate in Model II, and group was added as a subject-level covariate in Model III. Neither of these variables improved the model, however introduction of the interaction term between group and time significantly improved the model (LR = 15.88, *p* < .001) and yielded a significant main effect of time, and a significant effect of the group and time interaction (Model IV). As shown in Figure 3, subjects in the thin size condition showed a shift towards preferring thinner bodies from pre- to post-test, while subjects in the plus size condition showed the opposite pattern. Body size ideals in the plus size condition were significantly larger at post-test than those in the thin size condition (Figure 3). The model was further improved by adding sex (Model V: LR = 7.06, *df* = 2, *p* < .05) and location (Model VI: LR against Model V = 18.04, *df* = 2, *p* < .001) of subjects as both main effects and interaction terms with group and time, and showed a significant effect of location, such that subjects in the first village preferred thinner bodies in general than those in the second village. Addition of further potential confounding variables (age, BMI, dieting, education, time since last meal, and travel), did not improve the model any further (*p*s for likelihood ratios against Model V all > .1), and yielded no other main effects. Furthermore, none of the three-way interaction terms were significant (see full results in Extended data). As such, the interaction between time and group remained unmoderated by any of the potential confounders measured.

**Table 2:** Multilevel modelling analysis of predictors of ideal body size selected for each starter body: Models I-VI.

|  | I |  | II |  | III |  | IV |  | V |  | VI |  |
| --- | --- | --- | --- | --- | --- | --- | --- | --- | --- | --- | --- | --- |
| | $\beta$ | SE | $\beta$ | SE | $\beta$ | SE | $\beta$ | SE | $\beta$ | SE | $\beta$ | SE |
| Constant | -9.03 | 1.59 | -8.83 | 1.68 | -10.76 | 2.32 | -9.19 | 2.39 | -11.62 | 2.86 | -7.58 | 3.15 |
| **Time** |  |  | -0.41 | 1.11 | -0.41 | 1.11 | **-3.56** | **1.56** | **-3.56** | **1.56** | **-3.56** | **1.56** |
| Group |  |  |  |  | 3.76 | 3.15 | 0.66 | 3.33 | 0.52 | 3.31 | 0.57 | 3.14 |
| **Time\*Group** |  |  |  |  |  |  | **6.21** | **2.19** | **8.49** | **2.70** | **10.21** | **3.12** |
| Sex |  |  |  |  |  |  |  |  | 4.87 | 3.21 | 4.00 | 3.06 |
| Sex*Time*Group |  |  |  |  |  |  |  |  | -4.32 | 2.99 | -4.62 | 2.99 |
| **Location** |  |  |  |  |  |  |  |  |  |  | **-7.69** | **3.06** |
| Location*Time*Group |  |  |  |  |  |  |  |  |  |  | -3.31 | 2.99 |
| Log-likelihood: | 4688.25 |  | 4688.11 |  | 4686.69 |  | 4678.75 |  | 4675.22 |  | 4666.20 |  |
| Likelihood ratio<br>vs previous<br>model |  |  | 0.28 |  | 2.84 |  | **15.88** |  | **7.06** |  | **18.04** |  |

*Note.* Figures in bold indicate $p < .05$ . Models VII to XII are in Extended data.

**Figure 3.**
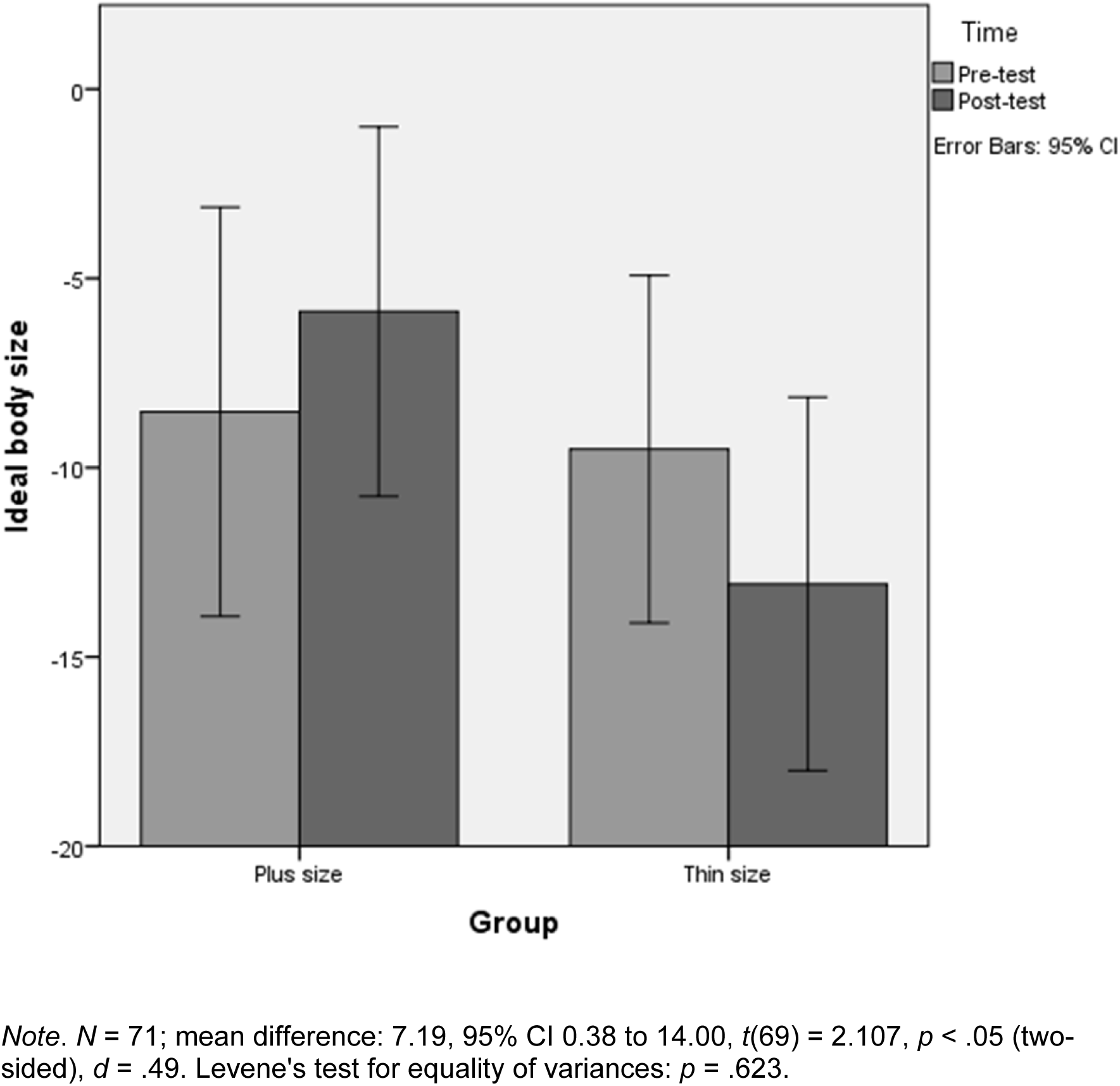
Pre-test to post-test difference in ideal body size between groups.

This study is the first to demonstrate experimentally that a short exposure to media images can change female body size ideals in a non-WEIRD population with very low to non-existent access to the media. Overall, just seeing a small selection of western media images of either thin or plus size female models induced change in subjects’ perceptions of ideal female body size in the direction of the images seen. These data are concordant with similar results in western samples^19-21^ and more importantly suggest that the capacity of visual experience to shape body ideals is not limited to subjects with a media-saturated environment. Furthermore, we have provided experimental evidence to support the assumption that the association between media access and both thinner body ideals^2^ and elevated body dissatisfaction^2,6^ in cross-cultural data, may be mediated in part by direct impacts of said media on internalised body ideals. The fact that none of our potential confounding variables moderated the experimental interaction, suggests that the impact of visual media on short term body ideals is not reliant on, for instance, recent prior media exposure or interest in weight loss.

It should be noted that, as per much visual media, we presented ‘aspirational’^19^ stimuli (i.e., healthy, attractive models in high status clothes) in the weight manipulations, and encouraged subjects to focus on their attractiveness. Thus we cannot determine whether our results were driven by the positive weight associations presented, or merely the visual experience itself. Furthermore, we used only one type of media images and future research should certainly consider the experimental impact of viewing other types such as magazines or real televisual stimuli on subjects, and should also consider both male stimuli and additional impacts on body satisfaction. Finally, although our subjects had very low to non-existent access to media in their villages, and despite the fact that the average participant tested had not left their village for 234 days, most have experienced some television when visiting towns or cities in the past. In other words, although body ideals in low media villages in this location are considerably larger than in the west^7^, it is possible that even modest prior experience may modulate the impacts seen here. Thus replication with a completely media-naïve population (not only in terms of TV access at home, but without any access to the visual media at all, even when travelling) would be ideal, albeit extremely difficult.

Nevertheless, we have shown that introducing western media images in a non-WEIRD population with very low prior media exposure can have an impact on their female body size ideals in a matter of a single 15-minute exposure. These results are important if we consider that grid electricity is gradually being introduced in our study site, that the local populations will soon have full, daily satellite TV access, and that TV consumption is associated with body dissatisfaction, low self-esteem, and disordered eating^4^. The globalization of the thin ideal in populations that are in the process of westernisation and/or modernisation^1^ is all the more worrying given that it usually coincides with rising levels of obesity, and therefore renders the dominant ideal of a slim or underweight female body even more difficult to attain. We thus strongly recommend further research on this pressing global challenge.

## METHODS

All study procedures and protocols were approved by Durham University’s Psychology Department Ethics Subcommittee (ref 10/09), and conformed to Durham University’s Data Protection Policy 2008 as well as to the ethical guidelines of the British Psychological Society.

### Study site

We selected two villages with no access to grid electricity and very low to non-existent media access on the remote Mosquito Coast of Nicaragua. The first village, San Vicente, was located in the Pearl Lagoon Basin and had no more than 40 adult inhabitants. The second village, Pedregal, was located on the nearby Patch River and had a maximum of 600 inhabitants. In San Vicente, one househould had a solar panel that was used for lighting, but there was no functional TV or access to other visual media in the village at the time of data collection. In Pedregal, a small number of households had recently acquired solar panels and satellite dishes and were therefore excluded from the study. Overall, the average hours of TV watched by the subjects in the last 7 days (including any TV watched while travelling) was 1.55 (mode: 0; median: 0). Further, the average subject had not left their village for 234 days. Together with participant observation, these statistics confirmed that the subjects had a very low exposure to visual media. More study site characteristics are in Table 1.

### Subjects

We tested 80 subjects (40 women) from 16 to 78 years (*M*= 30.4; *SD*=12.9) in total. Thirty-nine subjects were from San Vicente, where we tested all adults who were willing to participate and older than 16 years old. The remainder were recruited from Pedregal, where our rule was to test a similar number than in San Vicente and with an approximately equal number of men and women. All subjects provided consent and received $4 in local currency for their time. Subjects younger than 18 years old or older than 75 years old were tested with a parent or guardian present. Seven subjects from Pedregal who reported having access to satellite TV and watching it in the last 7 days, as well as 2 subjects who created the thinnest or fattest ideal body possible at both pre- and post-test (i.e. showed ceiling/floor effects), were discarded from analyses. Full sample characteristics are in Table 1.

### Materials and measures

For the ideal body task, we created 8 starter bodies in the software suite DAZ Studio 4.5. These bodies varied in shape (pear type vs. hourglass type), skin colour (light vs. dark), and clothing (swimsuit vs. low waist shorts and strapless bra), with each combination of shape/skin colour/clothing being represented. Importantly, the size of these starter bodies could be modified by the subjects (using left and right arrow keys over 20 increments) on a scale ranging from -20 (thinnest body) to +20 (heaviest body) and corresponding to a BMI range of 15 to 40. Half of the subjects received the first 4 bodies at pre-test, and the 4 remaining bodies at post-test, with the other half of the subjects receiving the opposite manipulation. Both at pre-test and post-test, the order of presentation of the bodies was randomised for each subject. Sizes selected for each body were recorded as the percentage of the screen width by which the mouse position deviated from the midpoint (-50% to 50%) and were treated as our dependent variable scores. Inspection of a Q-Q plot showed that these scores were approximately normally distributed, and multilevel model analyses were conducted in MLwiN 2.36.

For the manipulation, we used 144 photographs of either thin or plus size fashion models found on mainstream or specialist UK or U.S. retailers’ online catalogues or in popular women’s magazines. Although it was not always possible to determine which clothing size the models were wearing, most models in the thin and plus size condition ranged within UK sizes 4-6 and 16-28, respectively. All images showed women face-on, with a neutral background and with the full body visible (from head to knees at least), in non-sexually explicit poses. In each condition, there was an equal number of white and non-white ethnicity models, with two thirds of the models wearing clothes and one third wearing underwear or swimwear. The photographs were presented in pairs, giving 36 trials per condition, and the subjects were instructed to choose which of the two models they found more attractive. Within each pair, the models used looked similar and were matched for body shape, type of clothing, and pose. The number of same/different ethnic background pairs was counterbalanced both within and between experimental conditions. Further, the order of presentation of the pairs and the side of presentation of each photograph (left or right) were randomised for each subject.

All subjects also reported whether they had access to television (*in my house, in a neighbour’s house I visit, in a neighbour’s house I don’t visit, no TV in village*), and if they did, which type of television they had access to (*TV set-without satellite-and DVD player, Satellite TV, Satellite TV and DVD player, Other*), and how many hours they had watched it in the last 7 days. They also reported the last time that they travelled to other communities, how long they were away, and whether they had watched any TV while away (and how many hours in the last 7 days if they had). Subjects also reported how many hours ago they had last eaten and whether they were trying to change weight (*trying to lose weight, trying to increase weight, not trying to change weight*). Finally, we collected anthropometrics to compute the BMI and WHR of each subject. All sample characteristics are in Table 1.

### Procedure

The subjects were tested individually in a quiet room with a desk. The subjects already knew the experimenters from previous work in the area, and although they were aware of our main research interests (the electrification of the Mosquito Coast, and body ideals in Central America), they did not know the specific aim of the current experiment.

Upon providing oral informed consent, the subjects were assigned to condition (thin *vs.* plus size models) and presentation order of starter body quasi-randomly to ensure a similar number of subjects in each group (the experimenter was not blinded to group allocation). The subjects then completed the ideal body task on a laptop computer. Each body started at a random size, and the experimenter demonstrated to the subject how the size of the body could be changed using the left and right arrow keys, and showing the full possible range. The subjects were then instructed to modify each of the four bodies until these looked like their ‘ideal woman body’, meaning by this, ‘a woman body that to you is as attractive, good-looking, and sexy as possible’. They were also reminded several times that we were interested in their personal opinion and that they were no ‘right or wrong’ answers for this task. As most subjects were not used to computers, every effort was made to ensure that they knew how to use the left and right arrow keys and how this affected the size of the bodies. When the subject had finished a body, the experimenter asked again whether this represented their ideal female body before letting the subject proceed to the next body. When all 4 bodies were finished, the subject was told that ‘we will do something else now’, and received the manipulation. For each pair of photos, the experimenter asked the subject to choose their ‘favourite body’ between the two, or which one of the two bodies ‘looks better, or is a more attractive woman body in your opinion’. When the subject had chosen one body by tapping on it on the screen, the experimenter would display the next pair of photos by clicking the mouse, until all pairs had been displayed. This exposure to the thin/plus size models lasted approximately 15 minutes for each participant. The subjects then completed the ideal body task again, with exactly the same instructions as at pre-test (but with the 4 other starter bodies).

The session continued with an interview where subjects provided demographics and answered questions about media access, travel, etc. Finally, subjects’ height, weight, chest, waist, and hips were measured using an electronic scale and tape measure. These measurements were taken with the subjects dressed but without shoes. The subjects were given the opportunity to take their measurements themselves (with guidance), and anthropometrics for females were collected by a female experimenter or a female field assistant. A typical session lasted 40-45 minutes, and all subjects were interviewed in their native language (Creole English or Spanish).

## Acknowledgements

This research was supported by a Leverhulme trust project grant (RPG-2013-113) to LGB and MJT. We thank Robert A. Barton for helpful comments on the manuscript.

### Author contributions

Designed the study: JLJ with participation of all authors. Prepared the materials: MJT, JLJ, and LGB. Collected the data: JLJ and TT. Analysed the data: LGB an JLJ. Wrote the paper: JLJ and LGB with participation of MJT and TT.

### Competing financial interests

The authors declare that they have no competing financial interests.

### Materials & Correspondence

Correspondence and material requests should be addressed to.

### Data availability

The datasets generated during and/or analysed during the current study are available from the corresponding author on reasonable request.

---

Extended data. Multilevel modelling analysis of predictors of ideal body size selected for each starter body: Models VII-XII.

|  | VII |  | VIII |  | IX |  | X |  | XI |  | XII |  |
| --- | --- | --- | --- | --- | --- | --- | --- | --- | --- | --- | --- | --- |
| | $\beta$ | SE | $\beta$ | SE | $\beta$ | SE | $\beta$ | SE | $\beta$ | SE | $\beta$ | SE |
| Constant | -6.93 | 3.34 | -10.10 | 4.50 | -9.63 | 5.32 | -9.44 | 5.34 | -9.52 | 5.36 | -8.77 | 5.51 |
| **Time** | **-3.56** | **1.56** | **-3.56** | **1.56** | **-3.56** | **1.56** | **-3.56** | **1.56** | **-3.56** | **1.55** | **-3.56** | **1.55** |
| Group | 0.52 | 3.13 | 0.03 | 3.15 | 0.03 | 3.15 | 0.10 | 3.15 | 0.32 | 3.19 | 0.50 | 3.20 |
| **Time\*Group** | **10.90** | **3.29** | **14.38** | **4.57** | **14.10** | **5.37** | **14.17** | **5.38** | **13.95** | **5.39** | **14.54** | **5.64** |
| Sex | 4.16 | 3.06 | 3.98 | 3.05 | 3.93 | 3.06 | 3.84 | 3.07 | 3.63 | 3.11 | 2.94 | 3.33 |
| Sex*Time*Group | -4.11 | 3.07 | -3.94 | 3.07 | -3.88 | 3.17 | -3.80 | 3.17 | -3.36 | 3.22 | -4.18 | 3.76 |
| **Location** | **-6.77** | **3.45** | **-7.34** | **3.48** | **-7.45** | **3.54** | **-7.59** | **3.56** | **-7.88** | **3.62** | **-7.67** | **3.63** |
| Location*Time*Group | -2.37 | 3.33 | -2.33 | 3.33 | -2.27 | 3.37 | -2.70 | 3.44 | -1.77 | 3.63 | -1.82 | 3.64 |
| Travel | -1.83 | 3.21 | -1.13 | 3.26 | -0.98 | 3.39 | -1.07 | 3.39 | -0.91 | 3.42 | -1.06 | 3.41 |
| Travel*Time*Group | -2.31 | 3.41 | -1.57 | 3.58 | -1.67 | 3.74 | -1.70 | 3.74 | -2.37 | 3.84 | -2.18 | 3.85 |
| Age |  |  | 0.11 | 0.11 | 0.11 | 0.11 | 0.11 | 0.11 | 0.11 | 0.11 | 0.10 | 0.11 |
| Age*Time*Group |  |  | -0.12 | 0.12 | -0.13 | 0.12 | -0.13 | 0.12 | -0.11 | 0.12 | -0.11 | 0.12 |
| Last meal |  |  |  |  | -0.14 | 0.85 | -0.10 | 0.86 | 0.01 | 0.89 | 0.03 | 0.89 |
| Last meal*Time*Group |  |  |  |  | 0.09 | 0.90 | 0.27 | 0.95 | -0.08 | 1.05 | -0.02 | 1.06 |
| BMI |  |  |  |  |  |  | 0.00 | 0.01 | 0.00 | 0.01 | 0.00 | 0.01 |
| BMI*Time*Group |  |  |  |  |  |  | -0.01 | 0.01 | 0.00 | 0.01 | 0.00 | 0.01 |
| Education |  |  |  |  |  |  |  |  | -0.01 | 0.02 | -0.01 | 0.02 |
| Education*Time*Group |  |  |  |  |  |  |  |  | 0.01 | 0.02 | 0.01 | 0.02 |
| Dieting |  |  |  |  |  |  |  |  |  |  | -2.48 | 4.43 |
| Dieting*Time*Group |  |  |  |  |  |  |  |  |  |  | -1.70 | 4.66 |
| Log-likelihood: | 4665.18 |  | 4663.39 |  | 4663.36 |  | 4662.86 |  | 4662.13 |  | 4661.56 |  |
| Likelihood ratio vs previous model | 2.04 |  | 3.58 |  | 0.06 |  | 1 |  | 1.46 |  | 1.14 |  |

*Note.* Dieting and Travel were used as binary variables (i.e., is the subject trying to change their weight? *yes* or *no*; has the subject travelled in the last 7 days? *yes* or *no*). Ethnicity and earnings were not included because we considered that these were redundant with location and education, respectively. Further, separate analyses including ethnicity and earnings did not significantly change the results reported here.

## REFERENCES

1. Swami, V. Cultural influences on body size ideals: Unpacking the impact of westernization and modernization. European Psychologist 20, 44-51 (2015).

2. Swami, V. et al The attractive female body weight and female body dissatisfaction in 26 countries across 10 world regions: Results of the International Body Project I. Personality and Social Psychology Bulletin 36, 309-325 (2010).

3. Hogan, M. J. & Strasburger, V. C. Body image, eating disorders, and the media. Adolescent medicine: state of the art reviews 19, 521-546 (2008).

4. 4 Grabe, S., Ward, L. M. & Hyde, J. S. The role of the media in body image concerns among women: a meta-analysis of experimental and correlational studies. Psychological bulletin 134, 460-476 (2008).

5. Stice, E., Spangler, D. & Agras, W. S. Exposure to media-portrayed thin-ideal images adversely affects vulnerable girls: A longitudinal experiment. Journal of Social and Clinical Psychology 20, 270 (2001).

6. Becker, A. E., Burwell, R. A., Herzog, D. B., Hamburg, P. & Gilman, S. E. Eating behaviours and attitudes following prolonged exposure to television among ethnic Fijian adolescent girls. The British Journal of Psychiatry 180, 509-514 (2002).

7. Boothroyd, L. G. et al Television exposure predicts body size ideals in rural Nicaragua. British Journal of Psychology 107, 752-767 (2016).

8. Bearman, S. K., Presnell, K., Martinez, E. & Stice, E. The skinny on body dissatisfaction: A longitudinal study of adolescent girls and boys. Journal of youth and adolescence 35, 217-229 (2006).

9. Fallon, E. A., Harris, B. S. & Johnson, P. Prevalence of body dissatisfaction among a United States adult sample. Eating behaviors 15, 151-158 (2014).

10. Paxton, S. J., Neumark-Sztainer, D., Hannan, P. J. & Eisenberg, M. E. Body dissatisfaction prospectively predicts depressive mood and low self-esteem in adolescent girls and boys. Journal of clinical child and adolescent psychology 35, 539-549 (2006).

11. Johnson, F. & Wardle, J. Dietary restraint, body dissatisfaction, and psychological distress: a prospective analysis. Journal of abnormal psychology 114, 119 (2005).

12. Levine, M. P. & Harrison, K. in Handbook of eating disorders and obesity (ed J Kevin Thompson) 695-717 (2004).

13. Calogero, R. M., Boroughs, M. & Thompson, J. K. in The Body Beautiful (eds Viren Swami & A Furnham) 259-298 (Springer, 2007).

14. Anderson-Fye, E. P. & Becker, A. E. in Handbook of eating disorders and obesity (ed J Kevin Thompson) 565-589 (2004).

15. Swami, V. & Tovée, M. J. Perceptions of female body weight and shape among indigenous and urban Europeans. Scandinavian Journal of Psychology 48, 43-50 (2007).

16. Swami, V. & Tovée, M. J. Female physical attractiveness in Britain and Malaysia: A cross-cultural study. Body Image 2, 115-128 (2005).

17. Anderson, J. L., Crawford, C. B., Nadeau, J. & Lindberg, T. Was the Duchess of Windsor right? A cross-cultural review of the socioecology of ideals of female body shape. Ethology and Sociobiology 13, 197-227 (1992).

18. Henrich, J., Heine, S. J. & Norenzayan, A. The weirdest people in the world? Behavioral and brain sciences 33, 61-83 (2010).

19. Boothroyd, L. G., Tovée, M. J. & Pollet, T. V. Visual diet versus associative learning as mechanisms of change in body size preferences. PLoS One 7, e48691 (2012).

20. Robinson, E. & Christiansen, P. The changing face of obesity: exposure to and acceptance of obesity. Obesity 22, 1380-1386 (2014).

21. Robinson, E. & Kirkham, T. Is he a healthy weight? Exposure to obesity changes perception of the weight status of others. International Journal of Obesity 38, 663-667 (2014).

